# NATURAL VARIATION IN RADIOTOLERANCE IS ASSOCIATED WITH DIFFERENCES IN INNATE IMMUNE ACTIVITY AND S-PHASE TRANSCRIPTION IN DROSOPHILA MELANOGASTER

**DOI:** 10.64898/2026.05.07.723599

**Authors:** Llewellyn Green, Shahrzad Hajiarbabi, Erin Kelleher

**Author notes:** these authors contributed equally to this work (Llewellyn Green, Shahrzad Hajiarbabi).

## Abstract

Organismal tolerance of ionizing radiation is a complex trait whose genetic basis has been studied extensively, in large part due to its significance to human health and technological advancement. Conventional mutant screens in model organisms have revealed the paramount role of DNA damage response (DDR) and repair pathways in determining tolerance to ionizing radiation. However, uncovering natural genetic variation in radiotolerance is also of critical importance, as individual differences are associated with the differential susceptibility to cancer as well as differential response to radiation treatment. Genetic variation that underlies phenotype differences in natural populations often occurs in distinct genes and pathways as compared to the genes of major effect revealed by mutant screens, owing to the impact of natural selection on the former. We therefore sought to isolate natural variation in radiotolerance of *Drosophila melanogaster* by performing extreme QTL mapping. We generated a large genetically diverse multiparental population and exposed 3rd instar larvae to a semi-lethal dose or ionizing radiation. By sequencing surviving adults and comparing their haplotypes to unexposed controls from the same population, we identified a single major effect QTL spanning the 3rd chromosome centromere. The QTL contains 34 genes, none of which are previously implicated in radiotolerance. We interrogated the impact of these genes on radiotolerance through forward genetic analysis and RNA-seq. Our findings implicate diverse processes in radiotolerance including cell-cycle regulation and innate immune function.

## INTRODUCTION

Uncovering the genetic architecture of biologically significant traits is a major goal of any genetic analysis. However, there is a fundamental tension between the insights gained from classical genetic approaches (e.g. forward genetic screens and everything thereafter), and those gained from genetic mapping of natural variation. Forward genetic screens, by design, uncover genes and mutations with major effects on a phenotype of interest, whose biological functions are later revealed by detailed investigations of strong mutant phenotypes. By contrast, genetic mapping of natural variation generally shows that traits are highly polygenic, with individual variants having a modest or even undetectable effect on the phenotype (reviewed in Mackay and Anholt 2022). Furthermore, the effect of these variants on specific genes and biological processes are generally difficult to isolate. A major reason for this apparent contradiction is the action of purifying selection in natural populations, which removes deleterious alleles of major effect (Gazal et al. 2017; Zeng et al. 2018; O’Connor et al. 2019). Hence, variants that persist in natural populations have smaller effects on a more diverse array of biological processes.

Tolerance of ionizing radiation is a complex trait that has been studied extensively through classical genetic approaches, largely due to its significance to human health. Radiation exposure and sensitivity is often associated with cancer susceptibility, and radiation is a frontline therapy for many tumors (reviewed in El-Nachef et al. 2021). Extensive forward genetic screens in genetic model organisms such as yeast, *Caenorhabditis elegans a*nd *Drosophila melanogaster* have revealed that the ability to detect and repair DSBs is critical to radiotolerance, as these mutants often have the strongest radiosensitive phenotypes (Bennett et al. 2001; Boulton et al. 2002; Jaklevic and Su 2004). Similarly, rare recessive highly radiosensitive syndromes in humans are generally connected to homozygosity for mutations in DSB detection and repair factors such as *ATM* or *ligIV* (reviewed in El-Nachef et al. 2021). These genes are critical to recovery from radiation exposure because the major effect of radiation on cells is the production of DSBs both directly through striking DNA, and indirectly through the production of reactive oxygen species (ROS) in cells (reviewed in Griffin 2006).

In contrast, studies of natural genetic variation do not implicate DNA damage response and repair as the singular major determinant of individual differences in radiotolerance. A Genome Wide Association (GWAS) of radiotolerance in *D. melanogaster* did not observe that variants in DNA repair factors were enriched among the most significant SNPs (Vaisnav et al. 2014). Additionally, no SNPs in this study reached genome wide significance in association with radiotolerance, highlighting the persistent challenge of uncovering the genomic sources of natural variation. Similarly, a multitude of GWAS of radiation sensitivity among various categories of cancer patients generally reveal few (if any) significant SNPs, many of which do not obviously impact genes involved in DNA damage response (reviewed in Rosenstein et al. 2014; Naderi et al. 2022; Schack et al. 2022; Jandu et al. 2023). Rather, many of these SNPs are adjacent to genes involved in diverse biological processes including cell-cell adhesion, splicing, post translational modification and innate immune responses. The role of these processes in radiotolerance remain largely uncharacterized and poorly understood.

Quantitative trait locus (QTL) mapping with multi-parental panels (MPPs) is a powerful alternative to GWAS, which has proven success in uncovering complex genetic architectures in genetic models like *Drosophila, C. elegans* and mice (Churchill et al. 2004; King, Macdonald, et al. 2012; King, Merkes, et al. 2012; Snoek et al. 2019). In this approach, a small number of haplotypes are used as “founders” to generate a highly recombinant mapping population (reviewed in Long et al. 2014; Ladejobi et al. 2016). A major advantage of an MPP that all founder alleles are relatively common in the mapping population, meaning even rare variants confined to a single founder will be sampled many times. In contrast, rare variants will remain rare in a GWAS sample from a natural population, and their effects go undetected due to insufficient sampling. This represents a major limitation of GWAS based approaches, as rare genetic variants are more likely to exhibit phenotypic effects and are proposed to be a major source of phenotypic variation in natural populations (Pritchard 2001; reviewed in Mackay and Huang 2018).

Here, we mapped genetic variation in radiotolerance using the *Drosophila* Synthetic Population Resource, an MPP established from two groups of 8 founder haplotypes <u>(population</u> <u>B, King et al. 2012; King et al. 2012)</u>. We further took advantage of a bulk phenotyping approach, in which mixed populations were exposed to a semi-lethal dose of radiation, and survivors were sequenced to detect haplotypes in particular genomic regions that are associated with radiotolerance (Macdonald and Long 2022). We identified a single major effect QTL spanning the 3rd chromosome centromere, containing 34 genes, none of which are previously implicated in radiotolerance. We interrogated the impact of these genes on radiotolerance through forward genetic analysis and RNA-seq. Our findings implicate diverse processes including cell-cycle regulation and innate immune function in radiotolerance.

## METHODS

### Drosophila Stocks

Population B RILs, were obtained from Stuart Macdonald. Mutants and transgenic stocks for evaluation of candidate genes were obtained from the Bloomington, Vienna and Kyoto stock centers (Supplementary Table 1). In all experiments, stocks were maintained on standard cornmeal media at 22°C.

### Establishment and maintenance of the synthetic population

The synthetic population was established by pooling embryos from 703 Population B RILs. For each RIL, embryos were collected on yeast paste spread on agar grape plates and placed in separate standard cornmeal vials. Emerging adults were pooled across two large population cages (30×30cm) upon eclosion, which were maintained at census sizes of ∼1000 adults and fed standard cornmeal media. Generations were 2-3 weeks long and non-overlapping. Embryos were collected to establish the next generation by placing 4 bottles with media in each of the two cages. Bottles were randomly assigned to new cages to homogenize the population each generation.

#### Artificial Selection

In generations 6,7,9, and 10, 8 sets of 350-400 embryos were collected from population cages. Embryos were collected on grape-agar plates supplemented with yeast paste and placed on large petri dishes with standard cornmeal media. They were allowed to develop until the 3^rd^ instar (non-wandering stage) and then exposed to an acute dose of 40 Gy of ionizing radiation over ∼8.5 minutes (treated) or 0 Gy of ionizing radiation (control). Eclosed treated and control adults were frozen at –80°C until DNA extraction.

#### DNA extraction and sequencing

Genomic DNA was extracted from pools of frozen flies with Gentra Puregene Cell Kit protocol (Qiagen, 158767), including a few modifications described in Macdonald *et al*. (2022). PE-150 was performed on an Illumina NovaSeq 6000 S4 flow cell at 150PE at the University of Houston Sequencing Core, with coverage of aligned reads of 165X-500X coverage per sample. Pooled sequencing data for all samples can be found in the NCBI sequence read archive PRJNA1460791.

#### SNP calling and haplotype reconstruction

To estimate the frequency of 8 founder haplotypes in windows of 200 Kb and 500 Kb, we relied on the approach and scripts described in (Macdonald et al 2022). In brief, reads from our experimental pools, as well as published Illumina data from each of the 8 founders (King, Merkes, et al. 2012) were aligned to Release 6 of the *D. melanogaster* genome (dm6, Hoskins et al. 2015) using BWA (Li and Durbin 2009). bcftoolspileup, bcftoolscall and bcftoolsquery (Li 2011) were used to calculate the frequency of all SNPs not polymorphic in any founder in the experimental pools. LimsolveR (Van den Meersche et al. 2009; Soetaert et al. 2022 Nov 18) was then used to estimate the frequency of the 8 founder haplotypes for 200 Kb and 500 Kb windows, spaced 20 Kb apart.

#### QTL mapping

To isolate genomic windows in which founder haplotype frequencies differed depending on the treatment (irradiated vs control) we fit the following model to all haplotype windows for both window sizes (both 200Kb and 500Kb wide)

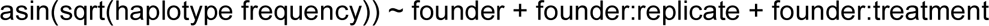

The LOD score was calculated from the *P*-value corresponding to the interaction between founder and treatment.

#### Identification of candidate variants in the QTL

Variants and corresponding genotypes for B6 founders were extracted from the *D. melanogaster* genome graph (Hickey et al. 2024) into VCF format using vg deconstruct (Liao et al. 2023). We focused on variants inside the QTL window where the sensitive haplotype (B6) differed from all population B founders, excluding low confidence variants where 3 or more founders could not be genotyped. Variants were classified as SNPs, small INDELS (100 nt or less), large INDELS (>1000 nucleotides) and complex (both alleles > 100 nt but neither is more than 10 fold larger than the other). Complex variants are difficult to characterize and were excluded from further analysis.

#### Impacts of candidate gene function on radiosensitivity

To evaluate the role of candidate genes in determining radiosensitivity we quantified adult eclosion for knock-down, mutant, and knock-up flies in the absence of radiation, and following an intermediate radiation dose (20 or 30 Gy) at the 3rd instar larval stage. Embryos or first instar larvae were collected from agar plates supplemented with yeast paste. Radiation (20 or 30 Gy) was administered to non-wandering third instar larvae along with matched unexposed controls. Pupae and eclosed adults were quantified for each of 3-6 replicate vials. The effect of genotype on adult eclosion following radiation treatment was assessed using logistic regression models as follows. For all models, vial was included as a random effect unless it resulted in overfitting of the model.

### For mutants

we compared heterozygous mutants for candidate genes and a control heterozygous mutant: *proboscipedia* (*pb[1] p[p]/TM3, Sb[1] Ser[1*] BDSC). We note that while *TM3* carries a loss-of-function mutation in *p53* <u>(Miller et al. 2016; Miller et al. 2020)</u>*, p53* mutations act recessively with regards to radiosensitivity <u>(Jaklevic and Su 2004; Sogame et al.</u> <u>2003)</u>, and there was no obvious difference between comparisons involving heterozygotes over TM3 (*FASN3, pb, RpL15, spok)* and heterozygotes over TM6 (*vtd* and *l(3)80Fj)*, which does not carry any p53 mutations. *pb[1]* is homozygous inviable arising from a developmental defect in the pupal stage (we did not observe *pb[1] p[p]* homozygotes in either our treated or control vials). Similarly we did not observe homozygotes of any of our candidate genes. The relationship between the candidate gene mutation and radiotolerance was therefore evaluated based on the significance of the interaction between treatment and genotype (candidate gene or *pb[1]*) in a logistic regression model where adult eclosion (or pupal lethality) was the response. In the case of *RpL15[301],* the very low survival rate following radiation treatment prohibited logistic regression. In this case we calculated the arcsin-transformed proportion of pupae eclosing in replicate vials and evaluated the difference between *RpL15[301]* and *pb[1]* using linear regression.

### For knock-down experiments with *daughterless*-Gal4 *(Da-Gal4)*

Knockdown genotypes containing a UAS-RNAi construct were compared to background matched controls (*Da-Gal4* and empty *attp2* or *attp40)* for adult eclosion or pupal lethality. The effect of knockdown on radiotolerance was evaluated based on the significance of the interaction between treatment and genotype (KD or *attp* background) in a logistic regression model where adult eclosion (or pupal lethality) was the response.

### For knock-down experiments with *actin*-Gal4 *(act-Gal4)*

knockdown genotypes containing *act-*Gal4 and UAS-RNAi construct were compared to balancer siblings (*CyO)* containing only the RNAi construct. Control crosses involving empty *attp2* and *attp40* were performed in parallel to rule out background effects. The effect of knockdown on radiotolerance was evaluated based on the significance of treatment in a logistic regression model where adult genotype (KD or CyO control).

### For CRISPR-activation experiments

knock-up genotypes containing *tub-Gal4, dCAS9* and target sgRNA were compared to balancer control siblings (*CyO:Tb*) that did not inherit *tub-Gal4* or *dCAS9.* Control crosses involving empty *attp2* were performed in parallel to rule out background effects. The relationship between knock-up and radiotolerance was determined from the significance of the interaction term between treatment and genotype (KU or balancer sibling) in the logistic regression model.

#### Sample Collection for RNA-seq

To identify pairs of background matched RILs with alternate haplotypes at the QTL peak, we took advantage of the published RIL genotypes (King, Merkes, et al. 2012). After identifying RILs that contain the B6 (sensitive) or B7 (tolerant) haplotype across the QTL window, we calculated the number of 10 KB windows where RIL pairs share the same founder haplotype. We selected two RIL pairs that share the same founder haplotype at the majority of windows outside the QTL (pair A: 57.3%, pair B: 64.4%).

Flies were maintained in small cages at 22°C, and embryos were collected on standard cornmeal media in small petri plates left in the cages overnight (35mm x 10mm). Embryos were allowed to develop for four days until the early third star wandering stage. For each RIL, 10-15 larvae were collected for three different treatment groups: 1) no radiation exposure, 2) 2 hours, and 3) 4 hours post radiation. The larvae were exposed to 30 Grays over 6 minutes and 40 seconds. Sampled larvae were homogenized in 200 μL TRIzol (Sigma-Aldrich) and kept at -80 °C. Three biological replicates for each of the two RIL pairs were collected on separate days. RNA extraction was performed using the TRIzol extraction following the manufacturer’s protocol (Thermofisher). RNA integrity number was determined by the University of Houston Sequencing and Gene Editing Core via Agilent 4200 TapeStation. RNA sequencing sample libraries were prepared using RNA Poly(A) selection and sequenced using the Illumina Novaseq X with the 1.5B flowcell by Texas A&M AgriLife Genomics & Bioinformatics Service Center. RNA-seq data for all samples can be found in the NCBI sequence read archive PRJNA1460791.

### RNA seq analysis

Annotated transcripts from the *D. melanogaster* reference genome (release 6.62) were quantified in the RNA-seq data using kallisto quant (Bray et al. 2016).

Differentially expressed genes between pairs, QTL haplotypes and treatment groups were identified using DESeq2 (Love et al. 2014). Prior to testing for differential expression we summed abundance across alternative transcripts of the same gene and then removed any genes with very low expression levels (<10 reads in at least >2 samples). To quantify TE expression, reads were mapped to the full dfam library of repeats for *D. melanogaster* including ancestral repeats and uncurated repeats <u>(Wheeler et al. 2013; Hubley et al. 2016; Storer et al.</u> <u>2021)</u>. To identify the differentially expressed genes and repeat families (DEGs) the following model was used ∼ Replicate:Pair + Pair + Haplotype + Treatment. We identified genes whose expression depended on an interaction between haplotype and pair using the model ∼ Replicate:Pair + Pair + Haplotype + Haplotype:Pair + Treatment. Log2FoldChange values and adjusted P values were obtained from the model for contrasts of interest (e.g. B6 vs B7). To visualize the shared and uniquely expressed genes between the datasets an upset plot was constructed using the UpsetR package in R <u>(Conway et al. 2017)</u>.

### Gene coexpression network construction

We identified modules of genes with highly correlated expression values using weighted gene co-expression network analysis <u>(WGCNA,</u> <u>Langfelder and Horvath 2008)</u>. Prior to WGCNA we accounted for batch effects (interaction of pair and replicate), and looked for hidden data structure (we found none) using SVA (Leek et al. 2012). We focused on modules that are positively co-expressed and detected correlation among genes using bi-midweight correlation. Modules with greater than 75% similarity were merged. The expression pattern of each module was summarized with a module eigengene for each sample, and correlated with explanatory variables using the following full model in limma (Ritchie et al. 2015):

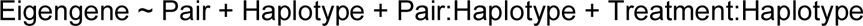

Effect estimates and *P-*values for individual terms were then estimated to detect relationships between modules and individual explanatory variables. Intramodular connectivity (kWithin) was calculated to identify highly connected genes within each module (Horvath and Dong 2008).

### Gene Ontology analysis

Gene Ontology (GO) enrichment analysis was performed to identify enriched biological processes among differentially expressed genes and genes within the WGCNA modules. GO enrichment analysis was conducted using the clusterProfiler package in R (Yu et al. 2012), with Benjamini-Hochberg (BH) correction for multiple testing and a q-value threshold of 0.05.

## RESULTS

### Extreme QTL Mapping

To isolate genomic regions associated with radiation tolerance we pursued extreme QTL mapping with population B of the *Drosophila* Synthetic Population Resource (Macdonald et al. 2022). We generated a large synthetic population from 703 RILs derived from 8 founder strains (B1-B7, and AB8). We maintained the population for 5 generations to allow for additional recombination. Starting with generation 6, we exposed non-wandering third instar larvae to a high dose of ionizing radiation (40 Gy), to enrich haplotype windows that confer radiation tolerance (Figure 1A). *Drosophila melanogaster* larvae are more radiosensitive than adults, as they are comprised of rapidly dividing mitotic cells (Paithankar et al. 2017). Our radiation dose yielded 83-88% mortality across replicate experiments (Figure 1B). Following next generation sequencing of pooled experimental and control samples (>165X for all samples), we uncovered 200 Kb genomic windows where the frequency of the founder haplotype was associated with radiation treatment (Figure 1C, Supplementary Table 2). The windows correspond to a single major effect QTL (LOD = 5.35) in the pericentromeric heterochromatin of chromosome 3 (3R:3127440-3527440).

**Figure 1.**
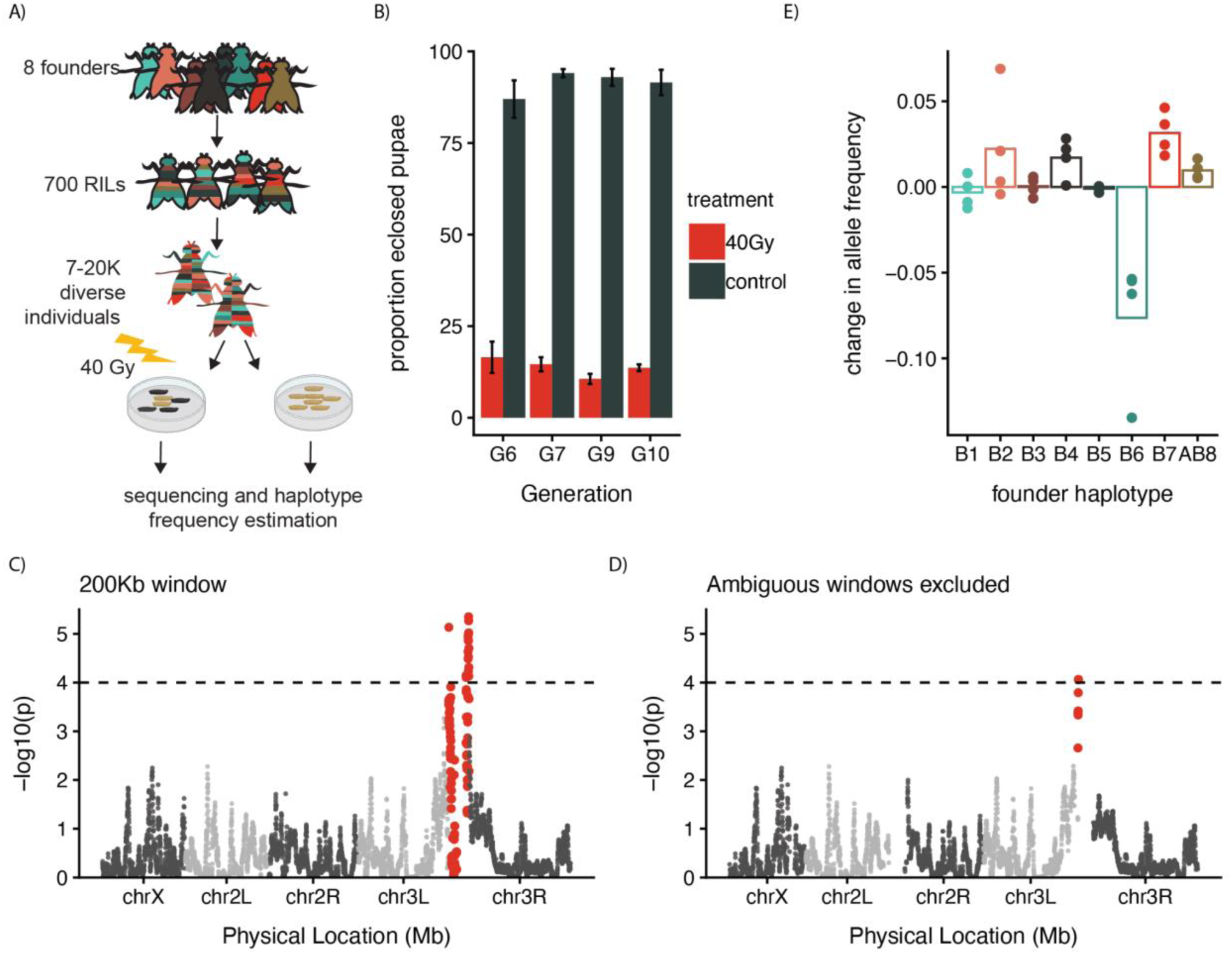
extreme QTL mapping of radiation sensitivity. A) Overview of extreme QTL mapping with the DSPR. B) Survival in replicates of treatment and control samples. C) LOD plot of haplotype association with radiosensitivity including all 200 Kb genomic windows. D) Change in allele frequencies of 8 founder alleles following radiation treatment at the LOD peak and adjacent left and right hand positions where the LOD scores are less than 1. Haplotypes frequencies are the same as in B. E) LOD plot of haplotype association with radiosensitivity for centromeric mapping, including a 5Kb window size in heterochromatic regions and excluding windows where some founder haplotypes cannot be resolved.

Haplotype inference close to the centromere is complicated by the high frequency of repetitive DNA (Hoskins et al. 2007), as well as the reduction in standing genetic variation when compared to euchromatic chromosome arms (Mackay et al. 2012). Indeed, for a standard 200 Kb window size we were unable to resolve the haplotype of all 8 founders for 17.5% of genomic windows (Supplementary Figure 1A), including all windows in the QTL peak. If anything, this ambiguity most likely decreases our power to detect QTL by creating uncertainty in the frequency of some haplotypes. Indeed, we observed that within our QTL, LOD scores drop as haplotype uncertainty increases while approaching the centromere (Supplementary Figure 1A-C). We reasoned that since recombination rates are low close to the centromere (Comeron et al. 2012), it was appropriate to employ a larger window size to estimate founder haplotype frequencies. Using a larger 500 Kb window, we identified 3 adjacent windows in the pericentromeric heterochromatin of chromosome 3R (positions 25676280,25696280, and 25716280) where all 8 founder haplotypes could be resolved and the LOD score exceeded the standard threshold of 4 (Figure 1D, Supplementary Figure 1B, Supplementary Table 3, (Linder et al. 2020; Macdonald et al. 2022)). The larger window otherwise had no obvious effect on the distribution of LOD scores throughout the genome (Supplementary Figure 1A-B). We conclude that there is at least one QTL that influences radiation sensitivity close to the 3rd chromosome centromere.

By examining haplotype frequencies in our irradiated samples when compared to controls, we observed that B2, B4 and B7 haplotypes moderately increased in frequency following radiation suggesting they are radiotolerant, while founder B6 strongly decreased in frequency, suggesting that it is radiosensitive (Figure 1E). We define the QTL window based on the genomic positions where the LOD scores are reduced by 3 when compared to the peak (-Δ3 LOD = 5.35 chr3L:25626280-chr3R:3667440, Figure 1D), as has been shown to be appropriately conservative for xQTL analyses with the DSPR <u>(Macdonald et al. 2022)</u>. While large, this region contains or partially overlaps with only 34 genes, of which 17are protein-coding genes and 17are non-coding RNAs (Supplementary Table 4). Several genes have interesting functions that are plausibly connected with radiation sensitivity including *Parp1*: (regulator of DNA repair), *verthandi* (DNA repair) and *Alg-2* (involved in apoptosis Figure 2A).

**Figure 2.**
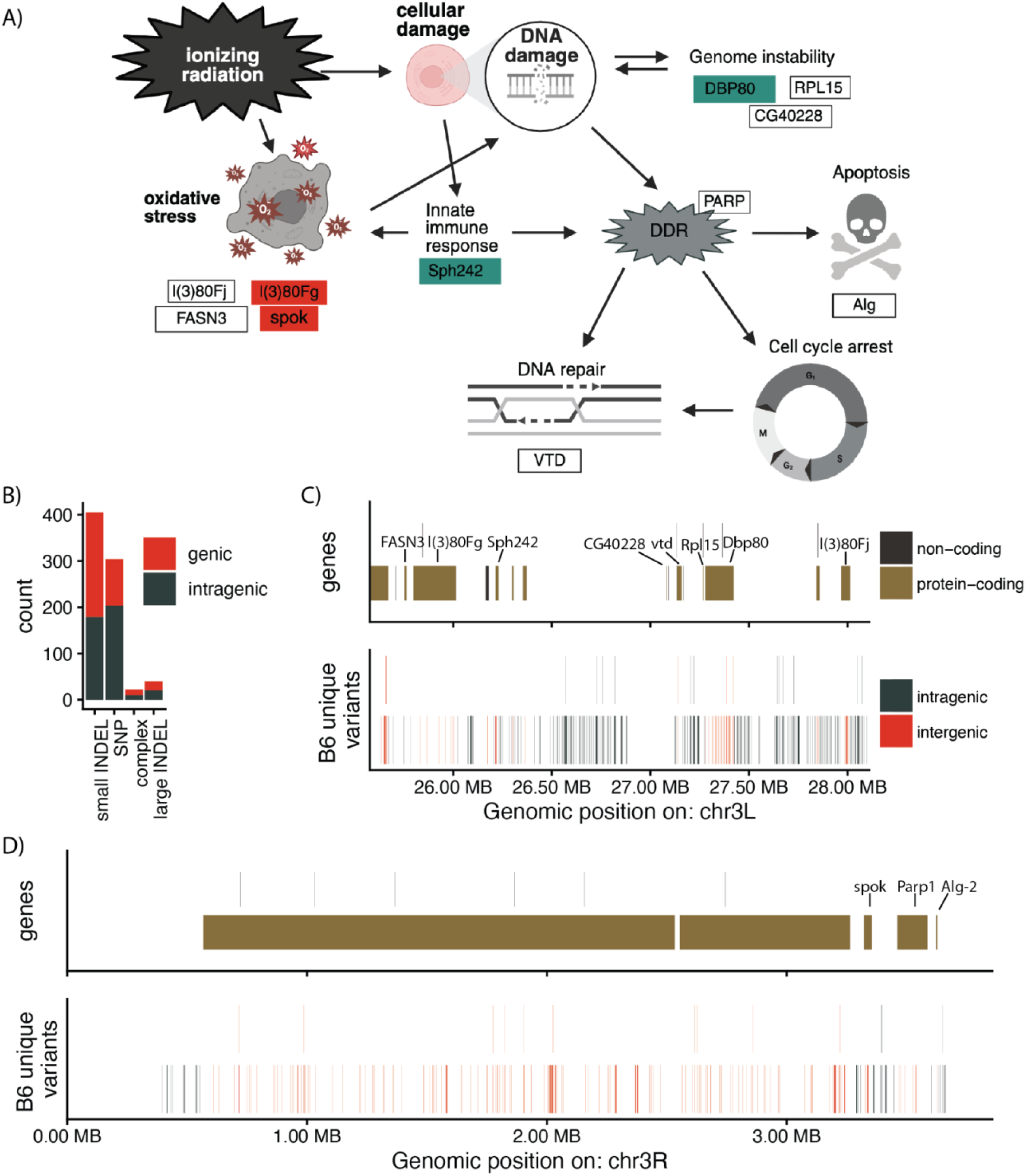
Candidate genes and variants within the QTL peak. A) Cartoon of the different potential function of candidate genes in the QTL peak. B) Frequency of different classes of molecular variants within the QTL with variants that are unique to B6, highlighting the proportion that are within 500 nt of an annotated gene. C and D) Browser view of the left (C) and right (D) hand side of the QTL peak, illustrating annotated genes (top panel) and molecular variants unique to B6 (bottom).

### Candidate causative variants

To identify potentially causative variants within the QTL, we compared the population B founder genomes using the *D. melanogaster* genome graph (Chakraborty et al. 2018; Chakraborty et al. 2019; Hickey et al. 2024). Given the extreme radiosensitivity of the B6 haplotype, we looked for variants that were unique to B6 among the population B founders. The genome graph contains 27,859 variants within the QTL, of which 777 could be genotyped in at least 5 population B founders and are unique to B6. These variants include 304 SNPs, 405 small INDELs (<100 nt), 22 large INDELs (>100 nt) and 40 “complex” variants that do not fit clearly in any of the aforementioned categories (Figure 2B). Manual inspection of the complex variants suggested that in most of these cases they include one or more large INDEL events along with accompanying INDELs or SNPS.

We next considered how the positions of B6-unique variants related to annotated genes within the QTL. In total, 363 variants occur within gene bodies or up to 2000 nt upstream of the transcription start site (TSS), potentially impacting the function of 18 genes in the QTL (Figure 2B-2C Supplementary table 4). The effects of these variants, if any, are most likely transcriptional or post-transcriptional, as no variants occur in an exon. In particular, 7 genes, including *eIF4B, Dbp80, l(3)80Fj, Sph242, CR14578, vtd* and *CR45181* have B6 unique variants within 2000nt (upstream or downstream) of the TSS, where cis-regulatory elements impacting promoter activity are most likely to reside <u>(Nègre et al. 2011)</u>.

### Forward genetics of candidate genes

The genetic basis of radiation tolerance has been studied extensively in *Drosophila* and other model organisms through forward genetic screens <u>(reviewed in</u> <u>Kelleher et al. 2025)</u>. Additionally, stereotyped biological responses to radiation have been curated in humans and other mammals from a healthcare perspective <u>(Vaiserman</u> <u>et al. 2018)</u>. Based on these two large bodies of literature, we identified 11 genes within the QTL region that could plausibly impact radiation sensitivity (Figure 2A). These genes have functions in the innate immune response *(Sph242)*, oxidative stress (*l(3)80Fg, l(3)80Fj, spok, FASN3),* genome stability (*Dbp80, Rpl15, CG40228)*, DNA damage response (*Parp1*), apoptosis (*Alg-2*), and DNA repair (*vtd,* Figure 2A). Of these 11, *Dbp80, l(3)80Fg, l(3)80Fj, Sph242, vtd, Parp1,* and *Alg-2* are also potentially impacted by variants that are unique to the B6 haplotype (Figure 2 C-D). We therefore evaluated the function of these candidate genes in radiotolerance using balanced mutants, RNAi knock-downs and CRISPR-activated overexpression (CRISPRa) (Ewen-Campen et al. 2017). We specifically quantified eclosion of pupae exposed to a high dose of radiation as non-wandering 3rd instar larvae, which was most comparable to the treatment in our original mapping (Supplementary Table 5).

We found weak evidence that *Dbp80* and *Parp1* promote survival following radiation exposure. Knock-down of *Dbp80* was associated with a reduction in eclosion rates when compared to irradiated balancer controls, however this was only significant for two of three RNAi constructs we examined (Figure 3A). Similarly, CRISPRa overexpression of *Parp1* was associated with increased eclosion following radiation treatment, but only when dCAS9 was inherited maternally (Figure 3D). Reciprocally, we did not observe that *Parp1* knock-down had a significant effect on radiotolerance (Figure 3A). However, it is important to note that *Parp1* expression depends on positive autoregulation that initiates in early embryos <u>(Tulin et al. 2002)</u>, which could render this gene refractory to RNAi and sensitive to maternal effects. We attempted to evaluate this through RT-PCR and were unable to do so due to the extremely low expression levels of *Parp1.* For both *Dbp80* and *Parp1* the subtle phenotypes associated with genetic manipulations suggest either that these genes play a minor role in radiotolerance or that their role is context-dependent.

**Figure 3.**
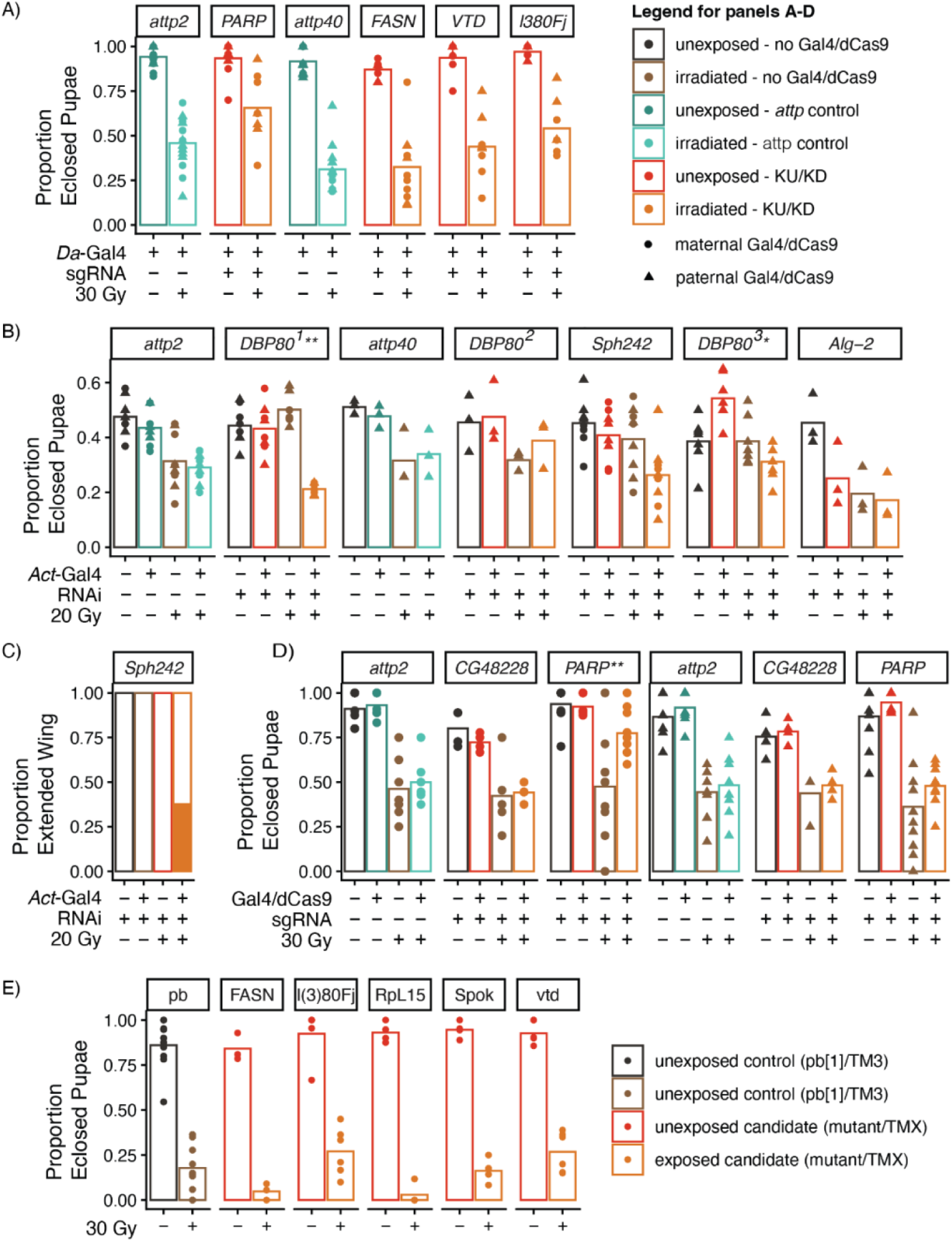
Forward genetics of candidate genes. The impact o1 candidate gene function on radiation tolerance in 3rd instar larvae was evaluated throug knock down (A-C), CRISPRa (D) and mutant alleles (E). Proportion of eclosed pupae (A,B,D,E) and proportion of eclosed adults with extended wings (C) are shown. Genes where altered expression or function had a significant effect on adult eclosion or wing extension following radiation exposure are indicated with asterisks. ** P > 0.01, *** P > 0.001.

We also observed an unusual phenotype following radiation exposure in *Sph242* knock-down flies. While adult eclosion following radiation did not differ from background matched or balancer controls (Figure 3B), ∼25% of eclosed adults show a wing extension phenotype that is associated with radiation exposure <u>(Seong et al. 2015,</u> Figure 3C). Hence, *Sph242* may be required to ensure proper development following radiation exposure.

### Expression differences between tolerant and sensitive haplotypes

In complement to forward genetic analysis of candidate genes, we employed RNA-seq to uncover differences in gene expression that may explain QTL haplotype-dependent differences in radiotolerance. For these experiments, we took advantage of the published RIL genotypes (King, Merkes, et al. 2012; King, Macdonald, et al. 2012) to identify nearly isogenic lines (NILs) which harbor different haplotypes within the QTL but share the same founder haplotype at the majority of windows outside of the QTL region. Since the B6 (sensitive) and B7 (tolerant) haplotypes showed the strongest opposing frequency changes in our original experiment (Figure 1C), we focused on comparing these haplotypes within our QTL, and identified two NILs that share 55% (pair A) and 62% (pair B) of founder haplotypes outside the QTL (Figure 4A). We then collected three replicates of third instar larvae from all four RILs for three different treatments: 1) unirradiated samples 2) 1 hour post-radiation, and 3) 4 hours post-radiation (30 Gy, Figure 4B). These timepoints were selected to represent important phases of DNA damage response. Cell cycle arrest is triggered approximately 1 hour post radiation to allow time for DNA repair (within 1-3 hours), and eventually apoptosis (around 4 hours) <u>(Dezzani et al. 1982; Jaklevic and Su 2004)</u>.

**Figure 4.**
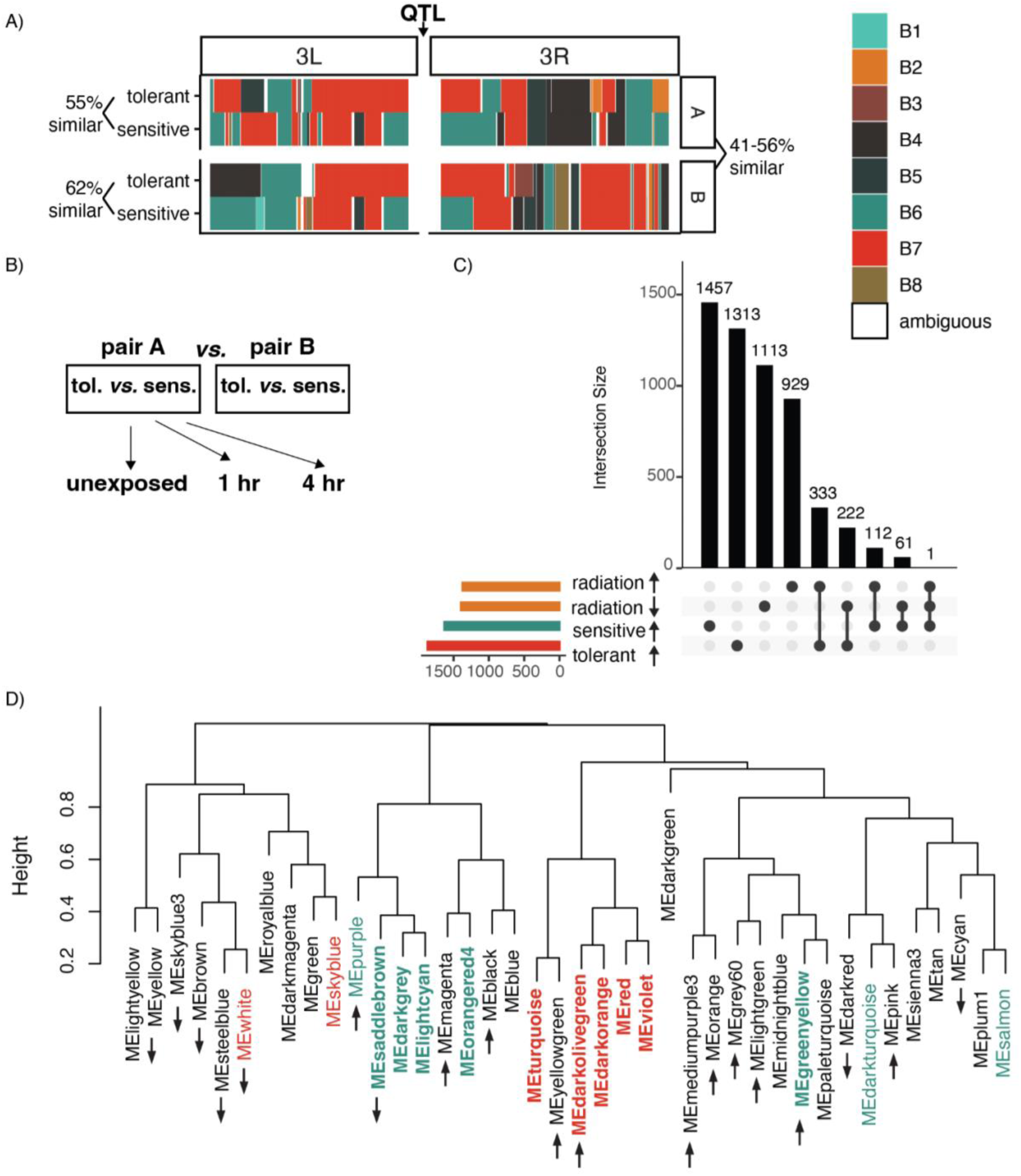
RNA-seq of radiation responses in sensitive and tolerant haplotypes. **A)** Haplotype map of the third chromosome, showing alternate alleles (B6 or B7) at the QTL peak and shared haplotypes outside of the QTL. Note that haplotype windows for Rlls do not extend into the pericentromeric heterochromatin where the boundaries of the QTL windows occur. B) Experimental design. C) Upset plot showing the total number and intersection of genes differentially expressed between haplotypes and following treatment **(1** or 4 hours post-radiation).**The** total number of upregulated or downregulated genes are indicated by horizontal bars, while the intersection of different groups of genes are indicated by vertical bars, connected by black dots below. D) Dendrogram of modules of positively co-expressed protein-coding genes revealed by WGCNA. Modules with increased expression in tolerant or sensitive haplotypes are colored red and green respectively. Modules in bold are not dependent on an interaction term between haplotype and RIL pair, or show a directionally consistent expression difference in both pairs. Increased and decreased expression post radiation are indicated by up and down arrows.

To reveal expression differences associated with radiosensitive and radiotolerant haplotypes, we first identified differentially expressed genes (DEGs). We identified 3499 genes differentially expressed between haplotypes, of which 1868 were upregulated in association with the tolerant haplotype, and 1631 are upregulated in the sensitive haplotype (Figure 4C, Supplementary Table 6). Among these are 9 genes in the QTL: 5 of which are more highly expressed by the tolerant haplotype (*Pzl, l(3)80Fg, spok, lovit, CR46250*) and 4 genes are more highly expressed by the sensitive haplotype (*Myo81F, Dbp80, Sph242, CR46123*, Supplementary Table 4). Two additional genes appeared differentially expressed however, on closer inspection they correspond to a single functional gene (*CG42598*) and an almost identical but likely non-functional paralog that is missing its 5’ UTR in the genome assembly (*CG41284*). The evidence for differential expression of these genes arises from a SNP in the first coding exon where the variant is shared between the functional and non-functional copies in the B7 but not the B6 haplotype.

All 7 differentially-expressed protein-coding genes are associated with a unique B6 variant within the gene body or up to 2000 nt upstream of the transcription start site, providing a potential mechanism for *cis-*regulatory variation (Figure 2B, Supplementary table X). Conversely, four of the differentially expressed genes (*Dbp80, Sph242, spok* and *CR46123*) show a significant interaction term between haplotype and NIL pair, indicating that the magnitude or direction of differential expression differs between Pair A and B (Supplementary table 4). This interaction reveals the effect of a *trans-*acting factor.

We also considered the relationship between QTL haplotype and radiation treatment. 729 genes that are differentially expressed between haplotypes also exhibit radio-responsive changes in expression (Figure 4C). However, none of these genes are located in the QTL. Furthermore, the expression response to radiation in these genes did not differ between haplotypes (i.e. the interaction term between haplotype and treatment was not significant for any gene). Collectively these results suggest that regulatory differences that may underlie the sensitive and tolerant phenotypes are constitutive, existing in both the presence and absence of radiation. We therefore focus on haplotype-dependent expression patterns in this manuscript, and will discuss radioresponsive expression changes more fully in a separate submission.

#### Identification of haplotype dependent modules of coexpressed genes

Given the large number of genes differentially expressed between haplotypes, we sought to group genes with related functions into co-regulated modules using Weighted Gene Co-expression Network Analysis (WGCNA, Langfelder and Horvath 2008). We confined this analysis to protein-coding genes, of which (>80%) have annotated biological functions in *D. melanogaster* <u>(Jenkins et al. 2022; Öztürk-Çolak et al. 2024)</u>. We identified 40 such modules of genes with positively correlated expression values (Figure 4D). Of these modules, 15 showed expression patterns that were associated with QTL haplotype (Supplementary Table 7). Of these 15 modules, 7 showed a significant interaction between RIL pair and QTL haplotype, indicating the impact of genetic variation outside the QTL (Supplementary Table 7). With regards to protein-coding genes in the QTL, 11 occur in modules with haplotype-dependent expression patterns (*l(3)80Fg, spok, lovit, eEIF4B, Alg-2, Parp1, Pzl, Sph242, Dbp80, Myo81F)*.

We also evaluated the relationship between module expression and radiation treatment. In total, 19 modules showed radiation-responsive changes in expression, of these 3 were also haplotype dependent: saddlebrown, darkolivegreen and greenyellow (Figure 4D). However no modules showed a haplotype-dependent radiation response (the interaction term between haplotype and treatment was not significant for any module). Thus, consistent with DESeq, the haplotype-dependent expression differences we uncovered through WGCNA were constitutive, existing in both the presence and absence of radiation.

#### Tolerant haplotypes exhibit enhanced innate immune activity

Since our goal in performing WGCNA was to better understand biological differences associated with QTL haplotype, we focused our attention on haplotype-dependent modules where the interaction with NIL pair was not significant (8 modules) or where the interaction was significant but the directional difference in expression was the same for both pairs (2 modules, Figure 4D, Figure 5A). To better understand the biological functions of these modules, we identified enriched GO terms among the genes within each module (Supplementary Table 8, Supplementary Table 9). We further identified the “hub” gene whose strong connectivity to other genes in the same module suggests its function may be fundamental to the network (Supplementary Table 10) (Carter et al. 2004; Carlson et al. 2006; Horvath and Dong 2008; Langfelder and Horvath 2008).

**Figure 5.**
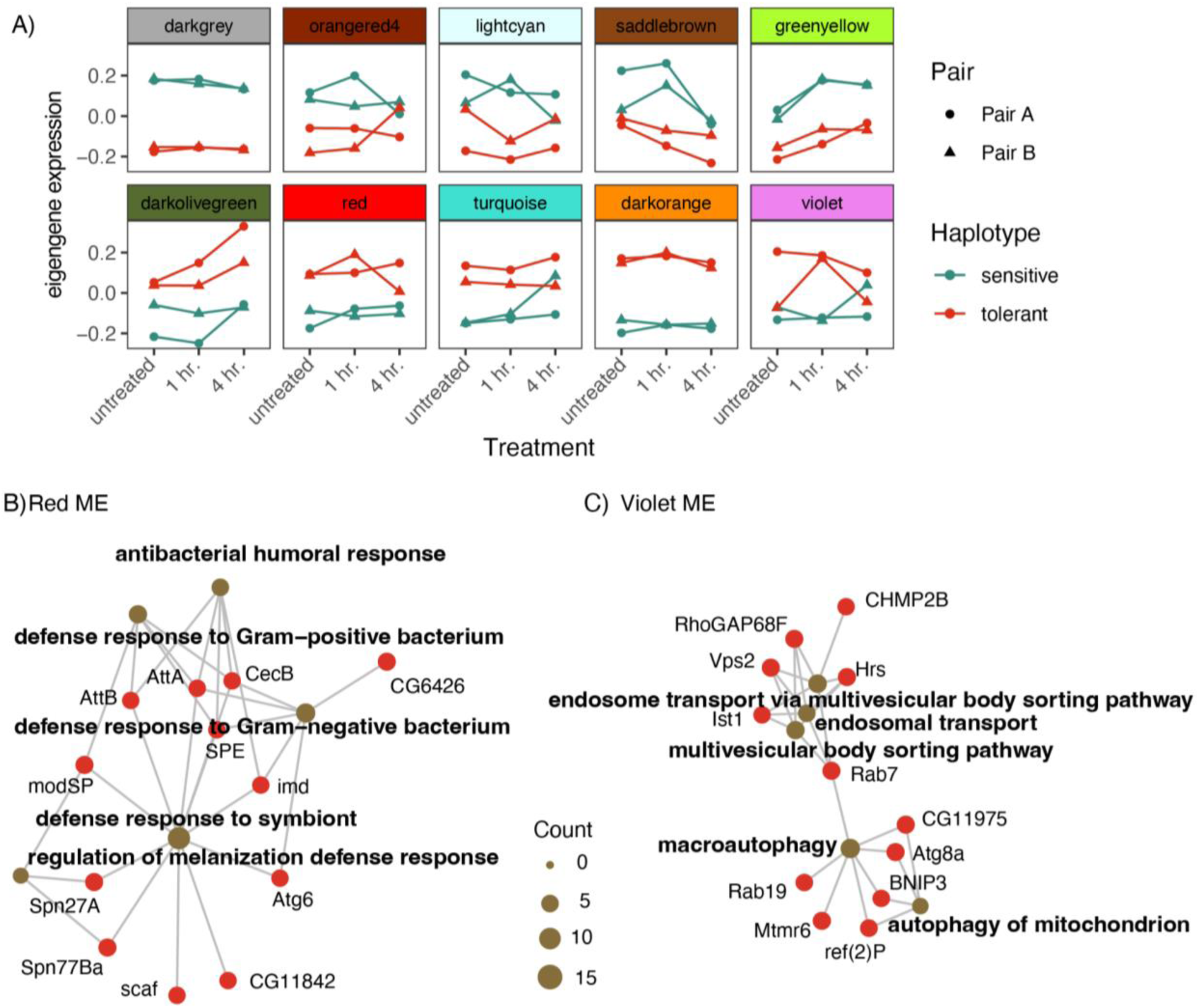
Haplotype dependent modules reveal increased innate immune activity in tolerant haplotypes. A) Eigenvalues for 10 modules showing haplotype-dependent expression patterns. Top row corresponds to modules with higher expression in sensitive haplotypes, while bottom row shows higher expression tolerant haplotypes. B-C) Cnet plots showing the top 5 most enriched GO Biological Process Terms for two modules with higher expression in tolerant: red (B) and violet (C).The subset of genes supporting the GO terms are shown as red circles, while the enriched GO term is represented as a tan circle. Size of the tan circle is dependent on the number of genes supporting each term.

Modules that are more highly expressed in tolerant RILs implicate a role for the innate immune system in radiotolerance. In particular, the red module contains multiple regulators of the antibacterial defense response including both upstream activators and downstream targets of the Toll and IMD pathways (Figure 5B). Similarly, the violet module is enriched for GO terms related to autophagy, an important innate immune response that clears and degrades damaged cells (reviewed in Kuo et al. 2018; Moreno-Blas et al. 2025). This implicates constitutive activity of innate immune pathways in *Drosophila* radiotolerance, consistent with observations in mammalian systems that Toll-pathway activation promotes radiotolerance <u>(reviewed in Liu et al.</u> <u>2018; Shi et al. 2017)</u>(Shi et al. 2017; Liu et al. 2018). Neither the red nor violet modules harbor genes located in the QTL, hence, the connection between innate immune activity and the QTL remains unclear.

The remaining three modules upregulated in tolerant are not clearly related to innate immunity or radiotolerance. Turquoise (contains *Pzl*) is enriched for GO terms related to spermatogenesis, which may imply more males, or males with larger testes, in those samples. Since males are generally more radiosensitive than females, this is unlikely to underlie an increase in radiotolerance (Parashar et al. 2008; Paithankar et al. 2017). In the case of darkolivegreen and darkorange (contains *spok* and *lovit*), no enriched GO terms are observed, however, the hub gene for the former is involved in tRNA synthesis.

#### Sensitive haplotypes show enriched transcription of S-phase transcripts

Modules upregulated in sensitive RILs, suggest increased DNA replication. In particular, saddlebrown and lightcyan (contains *eIF4B*) are enriched for genes involved in DNA replication and repair (Supplementary Tables 8,9). Similarly, darkgrey (contains *Myo81F*) is enriched for GO terms involved in nucleobase metabolism, which is fundamental to DNA synthesis (Supplementary Table Supplementary Tables 8,9). Transcription of many DNA replication and repair factors is activated by E2F1-Dp heterodimers, which are released from Rb at the start of S-phase (Duronio et al. 1996; Dimova et al. 2003; reviewed in Fischer and Müller 2017) (Figure 6A). E2F1-Dp also functions in a pathway that promotes apoptosis following exposure to ionizing radiation <u>(Zhou and Steller 2003; Wichmann et al. 2006; Wichmann et al. 2010)</u>. We therefore investigated whether E2F1-Dp and its known targets were enriched for genes upregulated in sensitive genotypes.

**Figure 6.**
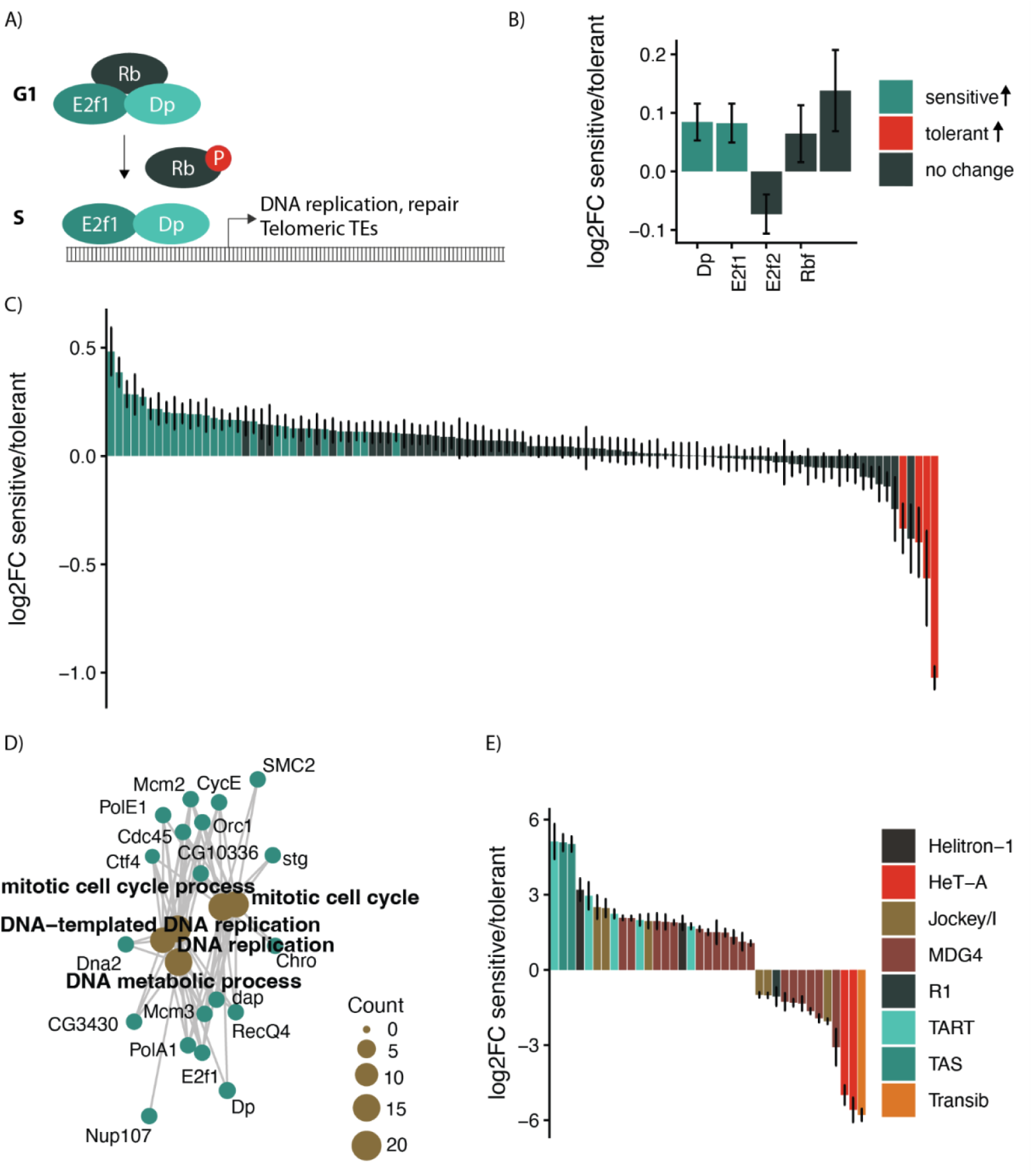
Sensitive haplotypes exhibit increased S-phase transcription. A) schematic of Rb, and E2F-Dp heterodimers regulate transcriptional changes at the start of 5-phase B) Expression differences in E2F/Dp/Rb transcripts between sensitive and tolerant haplotypes. C) Expression differences between sensitive and tolerant haplotypes in genes transcriptionally activated by E2Fl-Dp heterodimers from Dimova et al. 2003. D) Cnet plot of top 5 enriched GO terms (brown) among genes from panel C that are upregulated in sensitive haplotypes. Genes supporting the GO terms are shown as red circles, while the enriched GO term is represented as a tan circle. Size of the tan circle is dependent on the number of genes supporting each term. E) TE and repeat families differentially expressed between haplotypes, bars are colored according to TE or repeat family.

We observed that E2F1 and Dp are both modestly, but significantly, upregulated in sensitive genotypes, as are many of their known downstream positive regulatory targets (Figure 6B,C). Of 31 positive regulatory targets that are differentially expressed between the sensitive and tolerant haplotypes, 27 show higher expression in sensitive, indicating a strong bias towards increased expression in sensitive haplotypes (Sign-test *p= 3.4 x 10^-5^*). These genes are highly enriched for factors involved in DNA replication and repair (Figure 6D). We also observed that TART transposable elements and telomere associated sequence (TAS) repeats are upregulated in sensitive genotypes (Figure 6E), consistent with a recent study demonstrating that these sequences are positively regulated by E2F1-Dp (Liu et al. 2025). Finally the greenyellow module, which is increased in sensitive RILs as well as following radiation, is potentially connected to E2F1-DP. This module is enriched for genes involved in carbohydrate metabolism, which is known to be activated by E2F1-Dp in the polyploid fatbody tissues of the *Drosophila* larvae <u>(Zappia et al. 2021)</u>. Taken together, our data suggest more E2F1-Dp dependent gene activation in sensitive haplotypes.

## DISCUSSION

A major challenge in the study of natural variation in complex traits is the identification of the genetic variants and gene products that underlie differences between individuals (reviewed in Burgess 2022; Mackay and Anholt 2022; Long et al. 2025). A previous GWAS of natural variation in *D. melanogaster* radiotolerance illustrates this point, since heritability of radiotolerance was estimated as >80%, yet no significant SNPs could be associated with the phenotype <u>(Vaisnav et al. 2014)</u>. Here, using bulk-phenotyping of an MPP, we successfully uncovered a QTL for radiotolerance. Furthermore, while our study was well-powered in some respects (large number of RILs, good sequencing depth), it was limited in others (4 replicates or artificial selection and small pools of flies N=200). Hence, a more ambitious experiment could uncover additional radiotolerance QTL <u>(Macdonald et al. 2022)</u>.

Despite success in uncovering a QTL, we are not able to be conclusive about the causative gene or variant. While we found two genes within the QTL that have radioprotective functions and are differentially expressed between tolerant and sensitive haplotypes (*Dbp80,* and *Sph242)*, our data do not strongly support either gene as causative of the mapped QTL. Because these genes are radioprotective, their increased expression in sensitive haplotypes would be predicted to increase rather than decrease radiotolerance. Furthermore, none of these genes contain non-synonymous substitutions that are unique to the B6 sensitive haplotype, meaning that this inconsistency cannot be reconciled through the altered function of the encoded protein. Nevertheless, it is interesting to speculate that strong linkage disequilibrium within a pericentromeric QTL allows for genetic interactions between variants of multiple genes, which would be difficult to capture using standard forward genetics.

In light of the absence of a single strong candidate gene within our QTL, we examined gene expression differences associated with QTL haplotype that may underlie differences in radiotolerance. We cannot conclude that these expression differences are determinant of radiotolerance, however we uncovered two striking differences between tolerant and sensitive haplotypes that are potentially related to their disparate phenotypes. Tolerant haplotypes exhibit increased activity of innate immune pathways, while sensitive haplotypes show increased evidence of transcription of E2F-Dp1 S-phase targets. Both these patterns align with known determinants of radiotolerance.

A large body of research in mammalian systems implicates innate immunity in radiotolerance. In particular, Toll-like receptors are increased following radiation exposure in mammalian cells, and pharmacological activation of Toll-signaling promotes radiotolerance (reviewed in Liu et al. 2018; Shi et al. 2017; Burdelya et al. 2008). Two studies in *Drosophila* similarly suggest a function of innate immune pathways in post-radiation response <u>(Georgel et</u> <u>al. 2001; Sudmeier et al. 2015)</u>. Our work here suggests that genetic variation in innate immune function may also impact radiotolerance, and points to increased baseline innate immune activity as beneficial following radiation exposure. Our observations echo the recent discovery of a non-synonymous SNP in *STING1*, an important signaling factor in the mammalian interferon response, that is associated with differential toxicity following radiation exposure in head and neck cancer patients <u>(Naderi et al. 2022)</u>. It is interesting to speculate that the innate immune factors may contribute disproportionately to genetic variation in radiotolerance, as they are thought to be frequent targets or diversifying selection regimes imposed by pathogens <u>(Unckless et al. 2016; Minias and Vinkler 2022; Chapman et al. 2019)</u>.

A striking feature of the transcriptional profile of sensitive haplotypes is that there is enhanced transcription of S-phase genes, particularly the E2F1-Dp heterodimer and its known positive-regulatory targets <u>(Dimova et al. 2003)</u>. Since E2F1-Dp functions in a pathway that promotes apoptosis following exposure to ionizing radiation <u>(Zhou and Steller 2003; Wichmann et al.</u> <u>2006; Wichmann et al. 2010)</u>, increased E2F1-Dp could increase apoptosis rates in sensitive larvae following radiation exposure, thereby decreasing organismal survival. Alternatively, the relationship between E2F1-Dp and radiosensitivity could be related to S-phase directly. Dividing cells are generally more sensitive to DNA damage, owing to the need to repair and partition chromosomes into daughter cells <u>(Bergonie 1906; reviewed in Vogin and Foray 2013)</u>. In particular, radiosensitivity is high at the beginning of S-phase due to the low availability of homology dependent repair <u>(reviewed in Hustedt and Durocher 2016)</u>. Either way, our data suggest that genetic variation in highly conserved cell-cycle regulators could contribute to individual differences in radiotolerance.

Our work highlights that, while the challenge of identifying causative variants remains, QTL mapping from MPPs in model organisms is a powerful approach for uncovering natural genetic variation that goes undetected in GWAS of the same trait in the same species. Furthermore, while candidate genes and variants underlying QTL are often elusive, important biological insights can be gained through detailed investigations of mapped QTLhaplotypes. Innate immune responses and cell-cycle differences, which we implicate as sources of heritable variation in radiotolerance in *Drosophila* now provide testable hypotheses about how variation in such traits in human populations relates to radiation treatment and response.

## DATA AVAILABILITY

All DSPR RILs are available for purchase from the Bloomington Drosophila Stock Center (https://bdsc.indiana.edu/stocks/wt/dspr.html). The raw sequence data for pooled populations (experimental and control) as well as RNA-seq samples are available from the SRA archive (PRJNA1460791). Scripts used to reconstruct founder haplotype frequencies from pool seq data were previously described in Macdonald et. al (2022) and are available at https://github.com/sjmac/malathion-dspr-xqtl. Haplotype frequencies for 200 Kb and 500 Kb windows are available in Supplementary Table 2 and 3.

## ACKNOWLEDGEMENTS

We are grateful to Stuart Macdonald, Tony Long, and Richard Meisel for helpful discussion. We are also grateful to Sakina Motiwala and Lillian Pennington for assistance in maintaining the mapping population. This work was supported by R35GM138112 to E.S.K.

## REFERENCES

Bennett CB, Lewis LK, Karthikeyan G, Lobachev KS, Jin YH, Sterling JF, Snipe JR, Resnick MA. 2001. Genes required for ionizing radiation resistance in yeast. Nat Genet. 29(4):426–434.

Bergonie J. 1906. De quelques resultas de la radiotherapie et essai de fixation d’une technique radionnelle. C R Acad Sci. 143:983–995.

Boulton SJ, Gartner A, Reboul J, Vaglio P, Dyson N, Hill DE, Vidal M. 2002. Combined functional genomic maps of the C. elegans DNA damage response. Science. 295(5552):127–131.

Bray N, Pimentel H, Melsted P, Pachter L. 2016. Near-optimal RNA-Seq quantification with kallisto. Nat Biotechnol. 34:525–527.

Burgess DJ. 2022. Fine-mapping causal variants - why finding “the one” can be futile. Nat Rev Genet. 23(5):261.

Carlson MRJ, Zhang B, Fang Z, Mischel PS, Horvath S, Nelson SF. 2006. Gene connectivity, function, and sequence conservation: predictions from modular yeast co-expression networks. BMC Genomics. 7(1):40.

Carter SL, Brechbühler CM, Griffin M, Bond AT. 2004. Gene co-expression network topology provides a framework for molecular characterization of cellular state. Bioinformatics. 20(14):2242–2250.

Chakraborty M, Emerson JJ, Macdonald SJ, Long AD. 2019. Structural variants exhibit widespread allelic heterogeneity and shape variation in complex traits. Nat Commun. 10(1):4872.

Chakraborty M, VanKuren NW, Zhao R, Zhang X, Kalsow S, Emerson JJ. 2018. Hidden genetic variation shapes the structure of functional elements in Drosophila. Nat Genet. 50(1):20–25.

Chapman JR, Hill T, Unckless RL. 2019. Balancing selection drives the maintenance of genetic variation in Drosophila antimicrobial peptides. Genome Biol Evol. 11(9):2691–2701.

Churchill GA, Airey DC, Allayee H, Angel JM, Attie AD, Beatty J, Beavis WD, Belknap JK, Bennett B, Berrettini W, et al. 2004. The Collaborative Cross, a community resource for the genetic analysis of complex traits. Nat Genet. 36(11):1133–1137.

Comeron JM, Ratnappan R, Bailin S. 2012. The many landscapes of recombination in Drosophila melanogaster. PLoS Genet. 8(10):e1002905.

Dimova DK, Stevaux O, Frolov MV, Dyson NJ. 2003. Cell cycle-dependent and cell cycle-independent control of transcription by the Drosophila E2F/RB pathway. Genes Dev. 17(18):2308–2320.

Duronio RJ, Brook A, Dyson N, O’Farrell PH. 1996. E2F-induced S phase requires cyclin E. Genes Dev. 10(19):2505–2513.

Edgar BA, Orr-Weaver TL. 2001. Endoreplication cell cycles: more for less. Cell. 105(3):297–306.

El-Nachef L, Al-Choboq J, Restier-Verlet J, Granzotto A, Berthel E, Sonzogni L, Ferlazzo ML, Bouchet A, Leblond P, Combemale P, et al. 2021. Human Radiosensitivity and Radiosusceptibility: What Are the Differences? Int J Mol Sci. 22(13). doi:10.3390/ijms22137158. 10.3390/ijms22137158.

El Nachef L, Berthel E, Ferlazzo ML, Le Reun E, Al-Choboq J, Restier-Verlet J, Granzotto A, Sonzogni L, Bourguignon M, Foray N. 2022. Cancer and radiosensitivity syndromes: Is impaired nuclear ATM kinase activity the primum movens? Cancers (Basel). 14(24):6141.

Ewen-Campen B, Yang-Zhou D, Fernandes VR, González DP, Liu L-P, Tao R, Ren X, Sun J, Hu Y, Zirin J, et al. 2017. Optimized strategy for in vivo Cas9-activation in Drosophila. Proc Natl Acad Sci U S A. 114(35):9409–9414.

Fischer M, Müller GA. 2017. Cell cycle transcription control: DREAM/MuvB and RB-E2F complexes. Crit Rev Biochem Mol Biol. 52(6):638–662.

Gazal S, Finucane HK, Furlotte NA, Loh P-R, Palamara PF, Liu X, Schoech A, Bulik-Sullivan B, Neale BM, Gusev A, et al. 2017. Linkage disequilibrium-dependent architecture of human complex traits shows action of negative selection. Nat Genet. 49(10):1421–1427.

Georgel P, Naitza S, Kappler C, Ferrandon D, Zachary D, Swimmer C, Kopczynski C, Duyk G, Reichhart JM, Hoffmann JA. 2001. Drosophila immune deficiency (IMD) is a death domain protein that activates antibacterial defense and can promote apoptosis. Dev Cell. 1(4):503–514.

Griffin RJ. 2006. Radiobiology for the Radiologist. Elsevier.

Hendry JH, Simon SL, Wojcik A, Sohrabi M, Burkart W, Cardis E, Laurier D, Tirmarche M, Hayata I. 2009. Human exposure to high natural background radiation: what can it teach us about radiation risks? J Radiol Prot. 29(2A):A29–42.

Hickey G, Monlong J, Ebler J, Novak AM, Eizenga JM, Gao Y, Human Pangenome Reference Consortium, Marschall T, Li H, Paten B. 2024. Pangenome graph construction from genome alignments with Minigraph-Cactus. Nat Biotechnol. 42(4):663–673.

Horvath S, Dong J. 2008. Geometric Interpretation of Gene Coexpression Network Analysis. Miyano S, editor. PLoS Comput Biol. 4(8):e1000117.

Hoskins RA, Carlson JW, Kennedy C, Acevedo D, Evans-Holm M, Frise E, Wan KH, Park S, Mendez-Lago M, Rossi F, et al. 2007. Sequence finishing and mapping of Drosophila melanogaster heterochromatin. Science. 316(5831):1625–1628.

Hoskins RA, Carlson JW, Wan KH, Park S, Mendez I, Galle SE, Booth BW, Pfeiffer BD, George RA, Svirskas R, et al. 2015. The Release 6 reference sequence of the Drosophila melanogaster genome. Genome Res. 25(3):445–458.

Hustedt N, Durocher D. 2016. The control of DNA repair by the cell cycle. Nat Cell Biol. 19(1):1–9.

Jaklevic BR, Su TT. 2004. Relative contribution of DNA repair, cell cycle checkpoints, and cell death to survival after DNA damage in Drosophila larvae. Curr Biol. 14(1):23–32.

Jandu HK, Veal CD, Fachal L, Luccarini C, Aguado-Barrera ME, Altabas M, Azria D, Baten A, Bourgier C, Bultijnck R, et al. 2023. Genome-wide association study of treatment-related toxicity two years following radiotherapy for breast cancer. Radiother Oncol. 187:109806.

Kelleher ES, Hajiarbabi S, Green L. 2025. Extraordinary variation in radiation tolerance: Mechanisms and evolution. J Hered. 116(6):715–725.

King EG, Macdonald SJ, Long AD. 2012. Properties and power of the Drosophila Synthetic Population Resource for the routine dissection of complex traits. Genetics. 191(3):935–949.

King EG, Merkes CM, McNeil CL, Hoofer SR, Sen S, Broman KW, Long AD, Macdonald SJ. 2012. Genetic dissection of a model complex trait using the Drosophila Synthetic Population Resource. Genome Res. 22(8):1558–1566.

Kuo C-J, Hansen M, Troemel E. 2018. Autophagy and innate immunity: Insights from invertebrate model organisms. Autophagy. 14(2):233–242.

Ladejobi O, Elderfield J, Gardner KA, Gaynor RC, Hickey J, Hibberd JM, Mackay IJ, Bentley AR. 2016. Maximizing the potential of multi-parental crop populations. Appl Transl Genom. 11:9–17.

Langfelder P, Horvath S. 2008. WGCNA: an R package for weighted correlation network analysis. BMC Bioinformatics. 9(1):559.

Leek JT, Johnson WE, Parker HS, Jaffe AE, Storey JD. 2012. The sva package for removing batch effects and other unwanted variation in high-throughput experiments. Bioinformatics. 28(6):882–883.

Liao W-W, Asri M, Ebler J, Doerr D, Haukness M, Hickey G, Lu S, Lucas JK, Monlong J, Abel HJ, et al. 2023. A draft human pangenome reference. Nature. 617(7960):312–324.

Li H. 2011. A statistical framework for SNP calling, mutation discovery, association mapping and population genetical parameter estimation from sequencing data. Bioinformatics. 27(21):2987–2993.

Li H, Durbin R. 2009. Fast and accurate short read alignment with Burrows-Wheeler transform. Bioinformatics. 25(14):1754–1760.

Linder RA, Majumder A, Chakraborty M, Long A. 2020. Two synthetic 18-way outcrossed populations of diploid budding yeast with utility for complex trait dissection. Genetics. 215(2):323–342.

Liu M, Xie X-J, Li X, Ren X, Sun JL, Lin Z, Hemba-Waduge R-U-S, Ji J-Y. 2025. Transcriptional coupling of telomeric retrotransposons with the cell cycle. Sci Adv. 11(1):eadr2299.

Liu Z, Lei X, Li X, Cai J-M, Gao F, Yang Y-Y. 2018. Toll-like receptors and radiation protection. Eur Rev Med Pharmacol Sci. 22(1):31–39.

Long AD, Macdonald SJ, King EG. 2014. Dissecting complex traits using the Drosophila Synthetic Population Resource. Trends Genet. 30(11):488–495.

Long E, Williams J, Zhang H, Choi J. 2025. An evolving understanding of multiple causal variants underlying genetic association signals. Am J Hum Genet. 112(4):741–750.

Love MI, Huber W, Anders S. 2014. Moderated estimation of fold change and dispersion for RNA-seq data with DESeq2. Genome Biol. 15(12):550.

Macdonald SJ, Cloud-Richardson KM, Sims-West DJ, Long AD. 2022. Powerful, efficient QTL mapping in Drosophila melanogaster using bulked phenotyping and pooled sequencing. Genetics. 220(3). doi:10.1093/genetics/iyab238. 10.1093/genetics/iyab238.

Macdonald SJ, Long AD. 2022. Discovery of malathion resistance QTL in Drosophila melanogaster using a bulked phenotyping approach. G3 (Bethesda). 12(12):jkac279.

Mackay TFC, Anholt RRH. 2022. Gregor Mendel’s legacy in quantitative genetics. PLoS Biol. 20(7):e3001692.

Mackay TFC, Huang W. 2018. Charting the genotype-phenotype map: lessons from the Drosophila melanogaster Genetic Reference Panel. Wiley Interdiscip Rev Dev Biol. 7(1). doi:10.1002/wdev.289. 10.1002/wdev.289.

Mackay TFC, Richards S, Stone EA, Barbadilla A, Ayroles JF, Zhu D, Casillas S, Han Y, Magwire MM, Cridland JM, et al. 2012. The Drosophila melanogaster Genetic Reference Panel. Nature. 482(7384):173–178.

Minias P, Vinkler M. 2022. Selection balancing at innate immune genes: Adaptive polymorphism maintenance in Toll-like receptors. Mol Biol Evol. 39(5):msac102.

Moreno-Blas D, Adell T, González-Estévez C. 2025. Autophagy in tissue repair and regeneration. Cells. 14(4):282.

Mousseau TA, Møller AP. 2014. Genetic and ecological studies of animals in Chernobyl and Fukushima. J Hered. 105(5):704–709.

Naderi E, Schack LMH, Welsh C, Sim AYL, Aguado-Barrera ME, Dudding T, Summersgil H, Martínez-Calvo L, Ong EHW, Odding Y, et al. 2022. Meta-GWAS identifies the heritability of acute radiation-induced toxicities in head and neck cancer. Radiother Oncol. 176:138–148.

O’Connor LJ, Schoech AP, Hormozdiari F, Gazal S, Patterson N, Price AL. 2019. Extreme polygenicity of complex traits is explained by negative selection. Am J Hum Genet. 105(3):456–476.

Paithankar JG, Deeksha K, Patil RK. 2017. Gamma radiation tolerance in different life stages of the fruit fly Drosophila melanogaster. Int J Radiat Biol. 93(4):440–448.

Parashar V, Frankel S, Lurie AG, Rogina B. 2008. The effects of age on radiation resistance and oxidative stress in adult Drosophila melanogaster. Radiat Res. 169(6):707–711.

Pritchard JK. 2001. Are rare variants responsible for susceptibility to complex diseases? Am J Hum Genet. 69(1):124–137.

Ritchie ME, Phipson B, Wu D, Hu Y, Law CW, Shi W, Smyth GK. 2015. limma powers differential expression analyses for RNA-sequencing and microarray studies. Nucleic Acids Res. 43(7):e47.

Rosenstein BS, West CM, Bentzen SM, Alsner J, Andreassen CN, Azria D, Barnett GC, Baumann M, Burnet N, Chang-Claude J, et al. 2014. Radiogenomics: radiobiology enters the era of big data and team science. Int J Radiat Oncol Biol Phys. 89(4):709–713.

Sauer K, Knoblich JA, Richardson H, Lehner CF. 1995. Distinct modes of cyclin E/cdc2c kinase regulation and S-phase control in mitotic and endoreduplication cycles of Drosophila embryogenesis. Genes Dev. 9(11):1327–1339.

Schack LMH, Naderi E, Fachal L, Dorling L, Luccarini C, Dunning AM, Head and Neck Group of the Radiogenomics Consortium, Danish Head and Neck Cancer Group (DAHANCA), Ong EHW, Chua MLK, et al. 2022. A genome-wide association study of radiotherapy induced toxicity in head and neck cancer patients identifies a susceptibility locus associated with mucositis. Br J Cancer. 126(7):1082–1090.

Schaue D, Micewicz ED, Ratikan JA, Xie MW, Cheng G, McBride WH. 2015. Radiation and inflammation. Semin Radiat Oncol. 25(1):4–10.

Seong KM, Yu M, Lee K-S, Park S, Jin YW, Min K-J. 2015. Curcumin mitigates accelerated aging after irradiation in Drosophila by reducing oxidative stress. Biomed Res Int. 2015:425380.

Shi T, Li L, Zhou G, Wang C, Chen X, Zhang R, Xu J, Lu X, Jiang H, Chen J. 2017. Toll-like receptor 5 agonist CBLB502 induces radioprotective effects in vitro. Acta Biochim Biophys Sin (Shanghai). 49(6):487–495.

Smith AV, Orr-Weaver TL. 1991. The regulation of the cell cycle during Drosophila embryogenesis: the transition to polyteny. Development. 112(4):997–1008.

Snoek BL, Volkers RJM, Nijveen H, Petersen C, Dirksen P, Sterken MG, Nakad R, Riksen JAG, Rosenstiel P, Stastna JJ, et al. 2019. A multi-parent recombinant inbred line population of C. elegans allows identification of novel QTLs for complex life history traits. BMC Biol. 17(1):24.

Soetaert K, van den Meersche K, van Oevelen D. 2022 Nov 18. Package limSolve: solving linear inverse models in R. [accessed 2026 Jan 28]. https://cran.r-project.org/web/packages/limSolve/vignettes/limSolve.html.

Spurgin LG, Richardson DS. 2010. How pathogens drive genetic diversity: MHC, mechanisms and misunderstandings. Proc Biol Sci. 277(1684):979–988.

Sudmeier LJ, Samudrala S-S, Howard SP, Ganetzky B. 2015. Persistent activation of the innate immune response in adult Drosophila following radiation exposure during larval development. G3 (Bethesda). 5(11):2299–2306.

Unckless RL, Howick VM, Lazzaro BP. 2016. Convergent balancing selection on an antimicrobial peptide in Drosophila. Curr Biol. 26(2):257–262.

Vaisnav M, Xing C, Ku H-C, Hwang D, Stojadinovic S, Pertsemlidis A, Abrams JM. 2014. Genome-wide association analysis of radiation resistance in Drosophila melanogaster. PLoS One. 9(8):e104858.

Van den Meersche K, Soetaert K, Van Oevelen D. 2009. xsample(): AnRFunction for Sampling Linear Inverse Problems. J Stat Softw. 30(Code Snippet 1):1–15.

Vogin G, Foray N. 2013. The law of Bergonié and Tribondeau: a nice formula for a first approximation. Int J Radiat Biol. 89(1):2–8.

Yu G, Wang L-G, Han Y, He Q-Y. 2012. clusterProfiler: an R package for comparing biological themes among gene clusters. OMICS. 16(5):284–287.

Zappia MP, Guarner A, Kellie-Smith N, Rogers A, Morris R, Nicolay B, Boukhali M, Haas W, Dyson NJ, Frolov MV. 2021. E2F/Dp inactivation in fat body cells triggers systemic metabolic changes. Elife. 10(e67753). doi:10.7554/eLife.67753. [accessed 2026 Feb 24]. 10.7554/eLife.67753.

Zeng J de Vlaming R, Wu Y, Robinson MR, Lloyd-Jones LR, Yengo L, Yap CX, Xue A, Sidorenko J, McRae AF, et al. 2018. Signatures of negative selection in the genetic architecture of human complex traits. Nat Genet. 50(5):746–753.

